# Rare genetic variants in the endocannabinoid system genes *CNR1* and *DAGLA* are associated with neurological phenotypes in humans

**DOI:** 10.1101/168435

**Authors:** Douglas R. Smith, Christine M. Stanley, Theodore Foss, Richard G. Boles, Kevin McKernan

## Abstract

Rare genetic variants in the core endocannabinoid system genes *CNR1*, *CNR2*, *DAGLA*, *MGLL* and *FAAH* were identified in molecular testing data from up to 6.032 patients with a broad spectrum of neurological disorders. The variants were evaluated for association with phenotypes similar to those observed in the orthologous gene knockouts in mice. Heterozygous rare coding variants in *CNR1*, which encodes the type 1 cannabinoid receptor (CB1), were found to be significantly associated with pain sensitivity (especially migraine), sleep and memory disorders - alone or in combination with anxiety - compared to a set of controls without such *CNR1* variants. Similarly, heterozygous rare variants in *DAGLA*, which encodes diacylglycerol lipase alpha, were found to be significantly associated with seizures and developmental disorders, including abnormalities of brain morphology, compared to controls. Rare variants in *MGLL*, *FAAH* and *CNR2* were not associated with any neurological phenotypes in the patients tested. Diacylglycerol lipase alpha synthesizes the endocannabinoid 2-AG in the brain, which interacts with CB1 receptors. The phenotypes associated with rare *CNR1* variants are reminiscent of those implicated in the theory of clinical endocannabinoid deficiency syndrome. The severe phenotypes associated with rare *DAGLA* variants underscore the critical role of rapid 2-AG synthesis and the endocannabinoid system in regulating neurological function and development. Mapping of the variants to the 3D structure of the type 1 cannabinoid receptor, or primary structure of diacylglycerol lipase alpha, reveals clustering of variants in certain structural regions and is consistent with impacts to function.

## Introduction

The endocannabinoid system (ECS) plays an important role in the regulation of neurological activity throughout the central and peripheral nervous system [1–3], as well as in the regulation of cell division, metabolic, and immune processes in a variety of other tissues [4–6]. The type 1 cannabinoid receptor (CB1), encoded by the *CNR1* gene, is a key component of the ECS. CB1 is the most abundant G-protein coupled receptor in the brain, and is present at high levels in the neocortex, hippocampus and cerebellum [7]. CB1 is activated by the natural endocannabinoid agonists N-arachidonoylethanolamine (AEA), 2-arachidonoyl glycerol (2-AG) and a variety of related compounds [8]. It also binds the phytocannabinoid Δ9-tetrahydrocannabinol (THC), and a wide variety of synthetic agonists and antagonists [9]. At the cellular level, CB1 is located primarily on presynaptic termini of GABAergic and glutamatergic neurons in the brain [10, 11], where it binds 2-AG released by postsynaptic termini to down-regulate neurotransmitter release [12].

The type II cannabinoid receptor (CB2), encoded by the *CNR2* gene, is expressed in cells of the immune system (recently reviewed by Turcotte et al [13]) and is strongly induced in activated microglia in the brain [14]. CB2 is activated by the endocannabinoids AEA and 2-AG, and by the phytocannabinoids THC and β–caryophyllene [13, 15].

2-AG is the most abundant endocannabinoid in the brain, and is generated through the action of diacylglycerol lipase α (encoded by the *DAGLA* gene) by hydrolysis of diacylglycerol [16]. 2-AG is a mediator of retrograde signaling to presynaptic CB1 receptors to regulate neurotransmitter release [12]. 2-AG acts over longer distances during early development due to low monoacylglycerol lipase levels to regulate neuronal development at axonal growth cones [17, 18]. 2-AG is degraded by the action of monoacylglycerol lipase (encoded by the *MGLL* gene). AEA is generated mainly through the activity of N-acyl phosphatidylethanolamine phospholipase D, encoded by the NAPEPLD gene, and is degraded mainly by fatty acid amide hydrolase, encoded by the *FAAH* gene [19, 20].

Common single nucleotide polymorphisms in or near *CNR1*, *CNR2*, *FAAH* and *MGLL* have been reported to be associated with a variety of clinical phenotypes in candidate gene association studies (substance abuse disorders, cardiovascular disease risk factors, irritable bowel syndrome, migraine, chronic pain and mood disorders) [21–30]. The effect sizes are generally small, however, and replication studies in larger independent cohorts have been met with mixed results [31–35]. In contrast, the impact of rare genetic variation in genes associated with the endocannabinoid system has not been studied systematically. There are several reports of pathogenic deletions and duplications involving the ECS genes *CNR1*, *CNR2*, *DAGLA*, *MGLL* and *FAAH* with associated developmental phenotypes in the Decipher [36] and Clinvar [37] databases. There is also a reported association of *DAGLA* duplications with spinocerebellar ataxia-20 in OMIM [38] (entry 608687). However, the size of those structural variants is generally large (24 Mb on average) and they all impact multiple genes.

In this study, we investigate the phenotypic impact of rare missense variants in the core ECS genes: *CNR1*, *CNR2*, *DAGLA*, *MGLL* and *FAAH*, which encode CB1, the type 2 cannabinoid receptor (CB2), diacylglycerol lipase alpha, monoglyceride lipase, and fatty acid amide hydrolase, respectively. Phenotypes for were selected for evaluation based on their presence in mouse knockout strains for each gene.

## Methods

Two diagnostic gene panels were used in this study (Table S1). These panels were designed to identify chromosomal alterations associated mitochondrial disorders (NucSEEK Comprehensive – 1207 genes including *CNR1*, *MGLL* and *FAAH*) and epilepsy (EpiSEEK Comprehensive – 471 genes including *CNR1*, *CNR2*, *DAGLA*, *MGLL* and *FAAH*). Genetic sequencing data was generated from over 6,000 individuals using one or both of those gene panels in our CLIA/CAP certified laboratory, and clinical diagnostic reports were created, as follows.

DNA was extracted from saliva or blood samples using Genfind v2 (Beckman Coulter). Sequence-ready libraries were prepared using HaloPlex® target capture kits (Agilent Technologies) in conjunction with DREAM PCR amplification [39, 40] to avoid sample contamination. Sequencing was performed on the Illumina MiSeq platform to generate paired 250 base reads at average coverage levels of 465 for NucSEEK and 565 for EpiSEEK. Sequence reads were trimmed to remove adapters and low quality bases using Cutadapt and FastQC before alignment to the hg19 (GRCh37) reference sequence using BWA-MEM [41]. Qualimap [42] and Picard were used to generate alignment QC metrics such as insert size distribution, mismatch rates, and GC bias. Variant calls were generated by an ensemble [43] approach using FreeBayes, Platypus [44] and the GATK [45] UnifiedGenotyper [46]. For annotation and clinical review, a custom platform integrating data from multiple tools and databases was used (Ensembl variant effect predictor [47], SnpEff [48], Human Phenotype Ontology (HPO) [49], OMIM [38], ClinVar [50], 1000 genomes [51], ExAc [52], and dbNSFP [53]). SmartPCA [54] was used to stratify patients into subgroups based on analysis of the variants with respect to 1000 Genomes data. These data were stored in a database and reviewed in the context of clinical phenotype data provided by the patient's physician by Variant Scientists, Genetic Counselors, and Laboratory Directors. A comprehensive, user-friendly report was then generated for the patient's physician including pathogenicity scores generated according to ACMG guidelines [55].

Retrospective, aggregate analysis of variants in the core ECS genes for this study utilized only the pre-existing clinical information provided at the time of testing. Individuals were excluded if they did not sign a consent form for research use of their clinical testing data, or if the available clinical information was absent or inadequate. Adequate clinical information included an informative clinical summary, a comprehensive evaluation, or a completed checklist of standardized terms by a treating physician. Clinical summaries were translated to HPO terms by genetic counselors using the PhenoTips [56] software tool. A limited data set, containing only the above-described information and no personal identifiers, was used for the present study. Families were not contacted in regards to the current study. Thus, per the Office of Human Research Protection of the U.S. National Institutes of Health, this study does not qualify as human subjects research, and informed consent is unnecessary. However, the study was approved by the Courtagen Life Sciences ethics committee.

Rare variants in *CNR1*, *CNR2*, *DAGLA*, *MGLL* and *FAAH* having an allele frequency of approximately 0.003 or less in the ExAC database were identified. A Pubmed search of the literature relating to mouse knockouts of each of the five genes was carried out, and a list of the phenotypes described in those studies was compiled (Table S2). The mouse phenotypes were translated into general terms that could be related to human phenotypes in our test database (Table S2). Comprehensive lists of HPO terms were generated from a query of all scored tests in our database in which the five ECS genes were present (Tables S3 and S4). The terms were evaluated to generate subsets of HPO and text terms corresponding to each general phenotype for use as database queries (Tables S3 and S4). Counts of subjects with individual or multiple neurological phenotypes were generated through text-based queries of summary clinical phenotype data associated with de-identified reports and stored in a database. The phenotypes were scored positive if any of the included terms matched one or more of the clinical terms provided by the patient's physician. Non-neurological phenotypes were not evaluated because they were poorly represented in the clinical database, as almost all of the patients were being tested for neurological disorders.

A set of non-carrier controls were selected for each gene. The total number of non-carriers was 3,777 for DAGLA and CNR2, and 5,979 for CNR1, FAAH and MGLL. To simplify the analysis, 950 controls were selected at random for CNR1 and DAGLA, and ~500 controls were selected at random for CNR2, FAAH and MGLL. If the rare variant data for a particular gene was derived from two gene panels (this was the case for *CNR1*, *FAAH*, and *MGLL*), a proportional number of controls were selected from subjects sequenced using each of those panels. Counts of subjects with individual or multiple phenotypes were generated through text-based queries of summary clinical phenotype data as described above. Two-tailed Fisher exact tests were performed to evaluate the possible association of phenotypes present in carriers with rare variants, compared to controls, using a tool available at vassarstats.net (clinical research calculator number 3). The resulting P-values were Bonferroni corrected by multiplying by the number of general phenotypes tested (corrected P-values >1 were scored as 1). No correction was made for the number of genes tested, since they are all from a single pathway.

## Results

The numbers of subjects harboring rare variants were 22 for *CNR1*, 11 for *CNR2*, 35 for *DAGLA*, 34 for *MGLL* and 53 for *FAAH.* Data on the rare variants detected, including genomic location, coding consequences, allele frequencies in the ExAC, GnomAD and CLS databases, Polyphen2 and SIFT scores, PhyloP conservation and ExAC gene constraint values are summarized in Table S5. The human clinical phenotypes and HPO terms associated with each of the mouse knockout phenotypic categories detected are also presented in Table S5.

The results of Fisher exact tests for phenotypes putatively associated rare variants in the genes *CNR2*, *MGLL* and *FAAH* did not reveal any significant association between rare variants and any phenotype or combination tested (Table S6). The results for *CNR1* and *DAGLA*, however, revealed several significant associations between the presence of heterozygous rare coding variants and neurological phenotypes as indicated in Table 1.

**Table 1.** Association of mouse knockout phenotypes with rare variants in CNR1 and DAGLA.

| Gene* | Phenotype | Cases | Controls | OR | P-value |
| --- | --- | --- | --- | --- | --- |
| *CNR1* | Pain sensitivity | 9/22 | 156/950 | 3.5 | 0.036 |
| *CNR1* | Anxiety | 7/22 | 122/950 | 3.2 | 0.131 |
| *CNR1* | Memory disorder | 3/22 | 1/950 | 150 | 0.0003 |
| *CNR1* | Sleep disorder | 6/22 | 33/950 | 10.4 | 0.001 |
| *CNR1* | Urinary frequency | 2/22 | 19/950 | 4.9 | 0.301 |
| *CNR1* | Seizures | 11/22 | 402/950 | 1.4 | 1.000 |
| *CNR1* | Hearing loss | 1/22 | 31/950 | 1.4 | 1.000 |
| *CNR1* | Pain sensitivity and anxiety | 5/22 | 47/950 | 5.7 | 0.030 |
| *CNR1* | Sleep disorder and anxiety | 4/22 | 8/950 | 26.2 | 0.001 |
| *CNR1* | Pain sensitivity and memory disorder | 3/22 | 1/950 | 150 | 0.000 |
| *CNR1* | Pain sensitivity, memory and sleep disorder | 3/22 | 1/950 | 150 | 0.000 |
| *CNR1* | Pain sensitivity, anxiety and memory disorder | 2/22 | 0/950 | N/A | 0.003 |
| *CNR1* | Pain sensitivity, anxiety, memory and sleep disorder | 2/22 | 0/950 | N/A | 0.003 |
| *CNR1* | Pain sensitivity and sleep disorder | 3/22 | 7/950 | 21.3 | 0.005 |
| *DAGLA* | Anxiety | 2/35 | 177/950 | 0.26 | 0.211 |
| *DAGLA* | Neurodevelopmental disorder | 27/35 | 448/950 | 3.78 | 0.001 |
| *DAGLA* | Anxiety and neurodevelopmental disorder | 2/35 | 98/950 | 0.53 | 1.000 |
| *DAGLA* | Seizures | 30/35 | 595/950 | 3.84 | 0.032 |
| *DAGLA* | Seizures and neurodevelopmental disorder | 22/35 | 332/950 | 3.15 | 0.003 |

\*For each gene, the phenotypes that were evaluated are listed along with the number of cases or controls in which that phenotype was present out of the total number. Also listed are the odds ratio (OR), and P-values corrected for multiple testing. Confidence intervals for the odds ratios are provided in an extended version of the table in the supporting information (Table S7).

The allele frequencies of the ECS gene variants in our database of genetic testing results (CLS database) were compared with those in the public reference databases ExAC and gnomAD [57]. Across all five genes, the rare variants had an average allele frequency of 0.0004 in ExAC and gnomAD. The rare variants were enriched in the CLS database compared to gnomAD by 4.5-fold for *CNR1* and 6.7-fold for *DAGLA* (not including variants with zero frequency gnomAD). Eight of the 24 *CNR1* variants, and three of the 36 *DAGLA* variants had zero frequency in gnomAD. The fraction of variants with zero frequency in gnomAD for the other genes was 1/10 for *CNR2*, 8/53 for *FAAH* and 3/40 for *MGLL.*

We looked for shared phenotypes amongst subjects carrying the same rare variant. Two of the *CNR1* variants were observed in multiple samples (p.Glu93Lys in 2 samples, and p.Ala419Glu in 5 samples) - Table S5. All subjects with p.Glu93Lys had seizures and developmental delay. Two of the subjects with p.Ala419Glu had the shared phenotypes: anxiety, sleep disorder, abnormality of the autonomic nervous system and a third had a sleep disorder combined with seizures. Those phenotypes were absent in the other subjects with p.Ala419Glu.

Four of the *DAGLA* variants were observed in multiple samples (p.His810Gln in 7 samples, p.Arg815His in 5 samples, p.Ala858Val in 2 samples, and p.Leu988Val in 3 samples) - Table S5. Three of the subjects with p.His810Gln had infantile spasms and two had focal seizures. 5/6 of the subjects with p.Arg815His had seizures and developmental disorders and two had a brain abnormality. The subjects with p.Ala858Val did not have shared phenotypes. All three subjects with p.Leu988Val had seizures; two had developmental delay and the third had a brain abnormality in addition. One subject (ECS158) had two *DAGLA* variants, p.Arg197Trp and p.Ter1043GlnextTer29; the chromosomal phasing of those variants is unknown.

Additional genes with variants that could contribute to the observed phenotypes in the subjects, based on clinical reports, are noted in Table S5.

The 3D x-ray crystal structure of CB1 was published recently by two groups [58, 59]. We mapped the position of the rare coding variants onto the structure. Of the variants that could be mapped, 6/7 were in the 7-transmembrane cannabinoid binding domain (Fig. 1). The other *CNR1* variants, located near the N- or C-terminus, could not be mapped since those regions were deleted from the crystalized proteins. A similar exercise was performed for *DAGLA* using the primary structure and domain organization of the enzyme as described by Reisenberg, et al. [60] and NCBI Reference Sequence: NP_006124.1, since a crystal structure is not available. Most of the *DAGLA* variants mapped to the C-terminal end of the protein, a region that contains many regulatory phosphorylation sites (Fig. 2).

**Figure 1.**
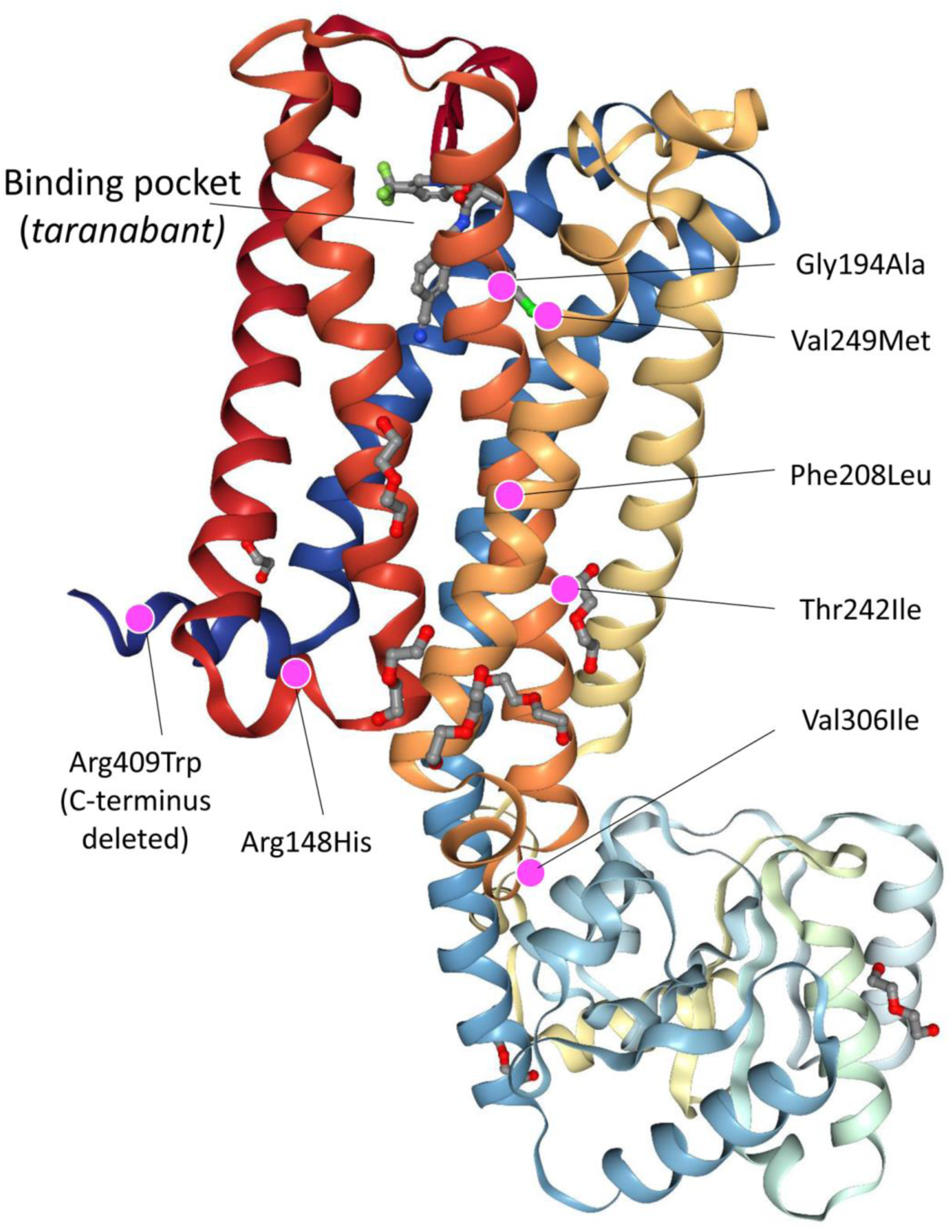
Depiction of the x-ray crystal structure of the type I cannabinoid receptor from PDB entry 5U09. The location of rare coding variants that could be mapped to the structure are shown by magenta dots, with annotations indicating the amino acid substitutions and the cannabinoid binding site.

**Figure 2.**
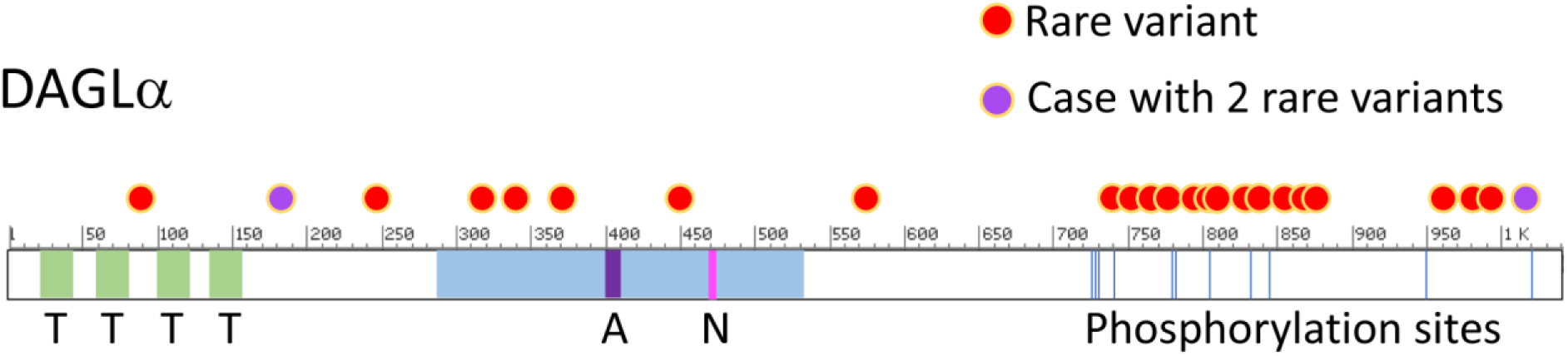
Mapping of rare variants to diacylglycerol lipase alpha. The location of rare coding variants relative to domains of the enzyme and regulatory phosphorylation sites are indicated, based on NCBI Reference Sequence: NP_006124.1. Red dot: rare variant; purple dots: two variants present in one subject; T (light green shading): transmembrane domain; light blue shading: lipase domain; A (purple shading): Active site; N (magenta shading): Nucleophilic elbow; blue vertical lines: phosphorylation sites.

## Discussion

We report a set of 155 rare missense or short deletion variants in the core ECS genes *CNR1*, *CNR2*, *DAGLA*, *FAAH* and *MGLL.* A subset of those variants, in *CNR1* and *DAGLA*, were found to be statistically associated with neurological phenotypes by case control analysis at the gene level. It is perhaps not surprising that *CNR2*, *FAAH* and *MGLL* were not associated with any phenotypes. Those genes are significantly more tolerant to missense and loss of function (LOF) variation than *CNR1* and *DAGLA*, based on ExAC gene constraint values [57, 61]. Furthermore, our testing focuses on neurological phenotypes, so metabolic or immunological phenotypes associated with those genes may not have been reported.

The *CNR1* variants were found to be associated with headache (including migraine), sleep and memory disorders, alone or in combination with anxiety, compared to a control set of 950 randomly selected patients without such *CNR1* variants. The Fisher odds ratios for the associated variants ranged from 4.5 up to 150, with the highest ratios being associated with sleep disorder plus headache (18.9), sleep disorder plus anxiety (26.2), memory disorder with or without headache (150) and the combined phenotypes of headache, memory disorder and anxiety and/or sleep disorder, which did not occur in any of the controls (Table 1).

Based on ExAC gene constraint values [57, 61], the *CNR1* gene is predicted to be somewhat tolerant of single missense and loss of function variants (missense z score = 2.7 and pLI = 0.15), but intolerant of dual heterozygous LOF variants (pRec = 0.78). All but one of the variants that could be mapped to the 3D structure of CB1 were in the 7-transmembrane domain. One additional variant, Ile339Val, would have mapped to the cytoplasmic domain if the region containing it had not been substituted in the 5U09 sequence. While we have no direct evidence that the variants we discovered impair function of the receptor, we note that three of the variants, p.Gly194Ala, p.Val249Met and p.Thr242Ile, map near the cannabinoid binding pocket in the 3-dimensional structure (Fig. 1). In addition, two other variants, p.Arg148His and p.Val306Ile, map near the boundary between the 7-transmembrane and cytoplasmic signaling domains. Such variants could affect signal transduction of the receptor. There were four variants (at residues 7, 27,57 and 93) that could not be mapped since the N-terminus was deleted from the structure. The N-terminal region is not required for cannabinoid binding, but may play a role in regulating the stability and surface expression of CB1[62]. We also note that the PolyPhen 2 deleteriousness scores of the *CNR1* variants are generally high (mean PP2 score = 0.85 with 11/24 ≥ 0.95).

Mice that are experimentally manipulated to harbor a non-functional *CNR1* gene (CB1 knockout), or that are treated with CB1 antagonists, display behaviors that suggest impacts related to pain sensitivity, anxiety, memory and sleep, amongst others (Table S2). The behaviors include elevated avoidance, freezing and risk-assessment behaviors [63], accelerated early learning and memory decline [64], food-seeking, and non-REM sleep alterations [65, 66]. There is a strong correlation between those phenotypes and those we observed in the present study (anxiety, sleep and memory disorders). Three of the eight subjects with rare *CNR1* variants and headaches in our study also have memory disorders (mainly short term memory), and two of those have comorbid anxiety and sleep disorders.

Two of the subjects with migraine headaches harbor variants in other genes that have previously been associated with migraine (*TRAP1* and *CACNA1A*) [67, 68]. However, the clinical correlation of the variants present in those subjects (*TRAP1* p.Ile253Val in subject ECS031 and *CACNA1A* p.Ser2423_Gly2424del in subject ECS100) with migraine and anxiety is uncertain. Interestingly, 2/5 subjects with a *CNR1* p.Ala419Glu variant (ExAC MAF = 0.00038) had the shared phenotypes: anxiety, sleep disorder, and abnormality of the autonomic nervous system. Only one of the other three subjects with that variant, however, had any of those phenotypes. These observations suggest that additional factors (genetic or environmental) may contribute to expression of the *CNR1*-associated phenotypes. Additional work will be required to resolve this.

The CB1 receptor encoded by *CNR1* binds the natural endocannabinoid ligands AEA and 2-AG. A theory of Clinical Endocannabinoid Deficiency (CED), first described by Russo, posits that migraine, fibromyalgia, irritable bowel syndrome and comorbid anxiety are related manifestations of reduced endocannabinoid tone, specifically AEA and 2-AG levels [69, 70]. These conditions frequently occur together, and a growing body of evidence suggests that they respond well to cannabinoid therapy, particularly THC [70]. Our results suggest that impaired CB1 signaling is associated with increased susceptibility to a related set of clinical phenotypes: migraine, sleep and memory disorders with co-morbid anxiety. It would be interesting to explore whether migraine, anxiety and sleep disorders in patients with rare deleterious variants in *CNR1* could be alleviated by treatment with THC or other CB1 agonists that may effectively stimulate the impaired and/or remaining functional CB1 receptors. In this regard, we note that a recent observational study of 121 adults with migraine reported a decrease in mean headache frequency from 10.4 to 4.6 headaches per month (p<0.0001) for patients receiving medicinal cannabis [71].

*DAGLA* encodes diacylglycerol lipase alpha, which is the major enzyme involved in 2-AG biosynthesis in the central nervous system. 2-AG levels have been reported to drop by ~80% in *DAGLA* knockout mice, resulting in pronounced anxiety and depression-like behaviors [72], and increased severity of kainate-induced seizures[73]. Developmental alterations observed in *DAGLA* knockout mice include altered cholinergic innervation of CA1 pyramidal cells of the hippocampus [18], and compromised adult neurogenesis in the hippocampus and subventricular zone [74]. In addition, altered axon formation in the midbrain-hindbrain region (associated with vision and locomotion) has been observed in response to transient morpholino *DAGLA* knockdown in zebrafish [75].

Heterozygous rare variants in *DAGLA* were found to be significantly associated with seizures, developmental disorders and abnormalities of brain morphology compared to controls. Based on ExAC gene constraint values, the *DAGLA* gene is predicted to be intolerant of missense and LOF variants (missense z score = 5.3 and pLI = 0.95 [57, 61]. We observed that 15/23 of the unique variants observed in our study map to the C-terminal third of the *DAGLA* coding sequence, a region that is predicted to contain numerous regulatory phosphorylation sites based on aggregate data from several biochemical studies [60]. This region appears to be more tolerant of variation than the catalytic domain, in which variation was sparse, despite its similar size. In contrast to mouse knockout models, anxiety phenotypes appear to be depleted in the clinical subjects - but the result was not significant. This raises the possibility that some of the variants might augment diacylglycerol lipase function. Further studies will be required to evaluate that possibility.

The frequency of seizures and developmental disorders in subjects with rare *DAGLA* variants were each elevated by a factor of approximately 4 compared to controls. The developmental disorders observed included several types of brain abnormalities: abnormalities of the cerebral white matter, brain malformations, posterior delayed myelination, heterotopia, and porencephaly with encephalomalacia. The prevalence of developmental disorders and brain abnormalities in these subjects aligns with the observations in animal models. Only two of the 35 samples with rare *DAGLA* variants harbored a likely pathogenic variant in another gene known to be associated with seizures and developmental delay (GRIN2A in subject 061 and MEF2C in subject 044). This represents about half the number of positive tests that would be expected based on the overall positive rate of the epiSEEK panel in our patient population (13%). Further work will be necessary to define whether rare variants in *DAGLA* are causatively associated with seizures and developmental disorders.

While the variants described in this study occur at extremely low frequencies in the general population, we suspect that patient populations with disorders of the nature studied here will be enriched for them. The allele frequencies observed in our test database for the rare *CNR1* and *DAGLA* variants were approximately 4.5-fold higher, on average, than those observed in the ExAC database. It will be interesting to evaluate what fraction of patients with migraine, with or without anxiety, sleep, or memory disorders, harbor low frequency or rare variants in *CNR1.* Similarly, the prevalence of rare *DAGLA* variants in patients with developmental disorders should be investigated.

## Acknowledgements

This study was supported by Courtagen Life Sciences, Inc. We thank Josee Dupuis for advice on the statistical analysis, and Ethan Russo and Anne Conyers-Hom for helpful comments on the manuscript.

## Supporting Table Captions

Table S1: Column 1 lists the genes contained in the EpiSEEK gene panel. Column 2 lists the genes contained in the NucSEEK gene panel.

Table S2: The phenotypes associated with knockout mice are listed under each gene in column 1. Column 2 gives the general phenotype that was used to select human phenotpes and HPO terms. Column 3 indicates whether the phenotypes are neurological. Column 4 provides PMIDs for corresponding mouse knockout publications.

Table S3: Column 1 provides a comprehensive listing of HPO terms derived from clinical reports for 5464 patients tested using Courtagen's NucSEEK and EpiSEEK gene panels. Column 2 provides the corresponding HPO descriptions. Columns 3-13 indicate the HPO terms corresponding to each of the general phenotypes derived from mouse knockouts according to the headers in row 3 (a “1” indicates that the term is included). Row 1 provides the strings of query terms used to evaluate the presence of general phenotypes in clinical reports of carrier and control subjects. Row 2 provides the frequencies of patients with the corresponding phenotypes amongst the 5464 patients.

Table S4: Column 1 provides a comprehensive listing of HPO terms derived from clinical reports for 3728 patients tested using Courtagen's EpiSEEK gene panel. Column 2 provides the corresponding HPO descriptions. Columns 3-8 indicate the HPO terms corresponding to each of the general phenotypes derived from mouse knockouts according to the headers in row 3 (a “1” indicates that the term is included). Row 1 provides the strings of query terms used to evaluate the presence of general phenotypes in clinical reports of carrier and control subjects. Row 2 provides the frequencies of patients with the corresponding phenotypes amongst the 5464 patients.

Table S5: Data for rare variants discovered in this study. Headers in row 1 are self explanatory.

Table S6: Association data for CNR2, FAAH and MGLL with column headers: Gene, Phenotypes that were evaluated, Number of cases or controls in which that phenotype was present out of the total number, odds ratio (OR), 95% confidence intervals for the odds ratios, uncorrected two-tailed P-values, and P-value corrected for multiple testing.

Table S7: This contains the same data as Table 1, with the addition of 95% confidence intervals for the odds ratios and the uncorrected two-tailed P-values. Columns identical to Table 1: Gene, Phenotypes that were evaluated, Number of cases or controls in which that phenotype was present out of the total number, odds ratio (OR), and P-value corrected for multiple testing.

